# Reversion to the Meta Body Form of Life after Complete Cell Death in Pure Liquid Water

**DOI:** 10.64898/2026.10.10.758155

**Authors:** Liaoqiong Fang, Yiting Wang, Hai Qiao, Jing Ma, Denggao Yu, Yongzhu Jiang, Hang Chen, Dan Song, Wenjing Song, Jin Bai, Qingyou Xia, Zhibiao Wang

## Abstract

The prevailing view holds that simpler animate beings predating the dawn of cellular life once existed, yet they have long since vanished from the present-day Earth. Our hypothesis posits that these animate beings persist to this day. To this end, we devised a long-term experiment spanning over a decade, keeping *Staphylococcus aureus* (*S. aureus*) within a sealed aqueous system of pure water. The findings illustrated that, upon the complete cell death (CCD) of the *S. aureus*, as cellular remnants progressively succumbed to dissolution, the aqueous solution underwent self-purification, bringing forth a class of lipid bilayer membrane microspheres containing RNA and proteins. These structures maintained long-term sustainable stability within the thermodynamic storm of water molecules and underwent dynamic size fluctuations over time – collectively exhibited phenomena indicative of being “alive”. This study reveals that the cell, upon complete cell death, undergoes reversion to a living entity simpler than the cell itself – the Meta Body (MB). The discovery of MB opens an entire re-conceptualization of what death ultimately brings.

## Introduction

The cell, in its profound intricacy and order, could never have emerged from sheer contingency. Hence, before the emergence of cellular life, primordial living entities must have existed. For the present study, we hypothesize that under certain specified conditions, cellular lives, upon complete cell death, undergo reversion to a simpler animate being—a more primordial form of life. Cells, placed within a sealed aqueous system of pure water (abbreviated as “sealed system”)—permitting thermal exchange with the external environment only, in time, shall yield to death and dissolution. If, after a sufficiently long duration of experimentation, the sealed system contains nothing more than a homogeneous solution of organic molecules, then the received wisdom—that simpler life forms predating cells have long since vanished from the Earth—would be axiomatically confirmed. Conversely, should the sealed system instead reveal structures that can respond to environmental changes, harness thermodynamic energy to sustain their own persistence, and undergo dynamic variations in size, then the inquiry into the origin of life would be endowed with both novel directions and unprecedented means.

The present work reports the findings of an 11-year sustained observational investigation.

## Methods and Materials

### 1. Bacterial culture

The *Staphylococcus aureus* strain ATCC 25923 (CICC 23656) was cultured on LB agar (HB0129; 40 g/L) and in LB broth (HB0128; 25 g/L). A single colony was picked from an LB agar plate and inoculated into 5–10 mL of LB broth, followed by overnight incubation (12–18 h) at 37 °C with shaking at 130 rpm. The overnight culture was then sub-cultured at a 1:100 dilution into fresh LB broth and grown under the same conditions to the exponential growth phase.

### 2. Preparation and storage of bacterial suspensions in sterile cryogenic vials

#### 2.1 Preparation of the initial bacterial suspension

The *Staphylococcus aureus* culture in the exponential growth phase was collected and centrifuged at 5,000 × g for 15 min at 4 °C (Eppendorf 5810R). The supernatant was discarded, and the pellet was collected, resuspended in sterile physiological saline and washed by centrifugation three times under the same conditions. Subsequently, an appropriate volume of sterile physiological saline (approximately 3 mL, negligible relative to the total suspension volume of 1000–1500 mL subsequently used for aliquoting) was added, and the pellet was gently pipetted and mixed to obtain the initial bacterial suspension.

#### 2.2 Preparation and aliquoting of bacterial suspensions in sterile deionized water

The initial bacterial suspension was diluted with sterile deionized water to the desired concentration. The adjusted suspension was aliquoted into sterile 2 mL cryogenic vials at 1.8 mL per vial. The caps were tightened, and the tube openings were further sealed with parafilm. Subsequently, every 50 vials were placed in a 100-well storage box at alternating positions, and the box was tightly wrapped in aluminum foil to protect from light. The boxes were then stored statically at room temperature or in incubators at specific temperatures according to the experimental requirements. All operations were performed under sterile conditions to avoid exogenous contamination. Each vial containing 1.8 mL of bacterial suspension and stored statically under light-protected conditions as described above was defined as a “sealed system.”

#### 2.3 Determination of the initial concentration of the packaged suspension and experimental grouping

The initial bacterial concentration of the sealed microtube system was finally determined by turbidimetry combined with the colony formation method. During dilution of the initial suspension with sterile deionized water, the concentration was estimated by turbidimetry using either a series of McFarland standards (prepared in 2014 for long-term observation) or spectrophotometric measurement of the absorbance at 600 nm (OD₆₀₀), and the dilution factor for the subsequent aliquoting was determined accordingly.

Immediately thereafter, an appropriate volume of the estimated suspension was subjected to tenfold serial dilution to prepare a dilution series of 10⁻²–10⁻⁷. One mL of each dilution was evenly spread onto the surface of agar plates. After the suspension was completely absorbed, the plates were inverted and incubated at 37 °C for 24 h, and colonies were then counted. Three parallel plates were prepared for each dilution, and the mean count was calculated (CFU/mL). The concentration of the original suspension was calculated according to the corresponding dilution factor, and the total bacterial count per vial was calculated based on 1.8 mL of suspension per vial (cells/vial).

Samples prepared in 2014 were divided into two groups:

(1) the room-temperature group for observing the time to complete bacterial death in the sealed microtube system, with an initial concentration of 2.55 × 10⁹ cells/vial;
(2) the long-term continuous observation group, with an initial concentration of 6.38 × 10⁹ cells/vial.

Samples for the short-term supplementary experiments were as follows:

(1) the room-temperature group, with an initial concentration of 1.91 × 10⁸ CFU/tube;
(2) the static 25 °C group for the complete bacterial death time experiment, with an initial concentration of 2.50 × 10⁸ CFU/tube;
(3) the static 37 °C constant group for the complete bacterial death time experiment, with an initial concentration of 3.02 × 10⁸ CFU/tube;
(4) the 37 °C constant-temperature group for observing the dynamic changes of particles in the sealed microtube system suspension, with an initial concentration of 2.50 × 10⁸ CFU/tube;
(5) the 37 °C constant-temperature purification experiment group, comprising two batches with initial concentrations of 3.02 × 10⁸ and 4.10 × 10⁸ CFU/tube, respectively.

### 3. Quantification of viable bacterial counts and determination of the time to total bacterial death

#### 3.1 Quantification of Viable Bacterial Counts

Sample tubes were randomly selected from the storage box. Based on the estimated viable count of 1 × 10⁴ CFU per sample, viable bacterial counting was performed using two procedures:

**(1)** **> 1 × 10⁴ CFU/sample** [1]: An appropriate aliquot of the bacterial suspension was subjected to tenfold serial dilution, using sterile deionized water as the diluent. One to three appropriate dilutions were selected, and 1 mL of each dilution was spread onto an LB agar plate, with one plate per dilution. The plates were first incubated at 37 °C for 1 day and then transferred to room temperature (∼25 °C) for a further 6 days of incubation. The viable count per sample was calculated from the colony counts on plates of each dilution and the corresponding dilution factor, according to the following formula:

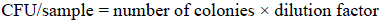
**(2)** **≤ 1 × 10⁴ CFU/sample**: The entire sample was divided equally into two portions, each of which was spread onto a separate LB agar plate. After the sample was removed, 1 mL of sterile PBS was added to the original vial, and the vial wall was washed by repeated pipetting for 10 s to detach the adherent bacteria; this wash solution was then spread onto a separate LB agar plate. All plates were incubated under the same conditions as described above (37 °C for 1 day, followed by 6 days at room temperature). Colonies on all plates were counted, and the total viable count per sample was calculated according to the following formula:

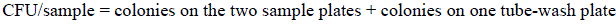

Three samples were taken for each time point.

#### 3.2 Determination of the time to complete cell death (CCD)

**(1)** **Room temperature group, 2.55 × 10⁹ cells/tube**: Aliquots of the *Staphylococcus aureus* suspension in cryogenic vials (1.8 mL per tube, initial bacterial count 2.55 × 10⁹ cells per vial) were stored at room temperature. During the 699-day experiment, samples were taken weekly during the early phase (days 0, 7, 14, 21, 28, 35, 42, 49, 56, 63, 70, 77, 84, and 91) and at monthly intervals during the later phase (days 151, 213, 274, 334, 393, 454, 515, 577, 638, and 699). At each time point, three vials were randomly selected as biological replicates, and viable bacterial counting was performed as described above. Survival curves were plotted with storage time on the x-axis and the number of surviving bacteria on the y-axis. Data are presented as mean ± standard deviation (SD).
**(2)** **Room temperature, 1.91 × 10⁸ cells/tube**: Aliquots of the *Staphylococcus aureus* suspension in cryogenic vials (1.8 mL per vial, initial bacterial count 1.91 × 10⁸ cells per vial) were stored at room temperature. Samples were taken at the following time points (days 0, 7, 14, 28, 42, 56, 84, 112, 140, 168, 196, 224, 252, 280, 308, 336, and 364). At each time point, three vials were randomly selected as biological replicates, and viable bacterial counting was performed as described above. Survival curves were plotted with storage time on the x-axis and the number of surviving bacteria on the y-axis. Data are presented as mean ± standard deviation (SD).
**(3)** **37 °C, 3.02 × 10⁸ cells/tube**: Aliquots of the *Staphylococcus aureus* suspension in cryogenic vials (1.8 mL per vial, initial bacterial count 3.02 × 10⁸ cells per vial) were stored at a constant temperature of 37 °C. Samples were taken at the following time points (days 0, 3, 6, 9, 12, 15, 18, 21, 24, 27, 30, 33, 36, 39, 46, 53, 60, and 67). At each time point, three vials were randomly selected as biological replicates, and viable bacterial counting was performed as described above. Survival curves were plotted with storage time on the x-axis and the number of surviving bacteria on the y-axis. Data are presented as mean ± standard deviation (SD).

#### 3.3. Define CCD and time to CCD

When no colony growth was observed on any of the parallel plates at a given time point (i.e., viable count = 0), sampling was continued at three additional independent time points using the same intervals as described above, with three vials per time point subjected to whole-sample plating. If no colony formation was observed in any of the samples from these three time points, the assay was terminated. The first time point at which a viable count of zero was detected was defined as the time of complete cell death, and the state of zero viable cells in this sealed microtube system was defined as complete cell death (CCD).

### 4. Nano-flow cytometry characterization of the microtube suspension

The concentration and size distribution of particles in the samples were analyzed using an Apogee A60-Micro nano-flow cytometer (Apogee Flow Systems, UK), which is equipped with 405 nm and 488 nm solid-state lasers. Prior to analysis, size calibration was performed using monodisperse silica nanoparticle standards (Apogee Mix #1493, refractive index 1.43, containing 180, 240, 300, 500, 590, 880, and 1300 nm particles), and instrument performance was verified using mixed reference microsphere standards. By adjusting the side-scatter threshold, the first gate was set above the noise background (i.e., the leftmost gate in the scatter plot) to eliminate electronic noise and optical background interference. Based on the size distribution of the standards, five additional size gates were defined: 80–180, 180–300, 300–590, 590–880 nm, and >880 nm. Sample vials were randomly selected for each time point, thoroughly mixed, and diluted with sterile deionized water to a particle concentration of 1 × 10⁶–1 × 10⁹ particles/mL. Data were acquired for 60 s at a flow rate of 1.5 µL/min with a total sample volume of 250–300 µL, and a solvent-only control was included to verify the reasonableness of the threshold settings. The accompanying software was used to analyze the event data: particle diameter was calculated from the 405 nm MALS signal, sample purity was assessed from the 488 nm Grn fluorescence signal, and the number of particles in each size interval was calculated (particles/vial). All experiments were independently repeated three times.

### 5. Detection of RNA and Protein

The membrane microspheres and the outer fluid were separated from the samples by ultrafiltration using an Amicon® Ultra-15 Centrifugal Filter Devices (100kDa, Millipore, UFC9100), centrifuged at 1000g for 10 minutes at room temperature. The membrane microspheres retained on the filter membrane were washed twice with PBS and then resuspended in PBS to a final volume of 200µL. Half of the suspension (100µL) was used for RNA detection, and the other half (100µL) was used for protein detection. Meanwhile, two equal aliquotes (100µL each) of the filtrated outer fluid were taken for RNA and protein detection, respectively.

Total RNA, including miRNA, was extracted from the samples using the miRNeasy Serum/Plasma Advanced Kit (QIAGEN, 217204) following the manufacturer’s protocol. Briefly, the samples were lysed with Buffer RPL and incubated at room temperature for 3 min. Proteins were precipitated by adding Buffer RPP, followed by vigorous vortexing and a further 3-min incubation at room temperature, and the mixture was then centrifuged at 12,000×g for 3 min. The resulting supernatant was mixed with an equal volume of isopropanol and applied to a RNeasy UCP MinElute spin column. After sequential washes with Buffer RWT, Buffer RPE, and 80% ethanol, the column was dried by centrifugation at full speed with the lid open for 5 min. Finally, RNA was eluted in 20 μl of RNase-free water, by incubating for 1 min followed by centrifuging for 1 min. RNA concentration was measured using a Qubit™ 4 fluorometer (Thermo Fisher Scientific), and RNA integrity and size distribution were assessed using a 5200 Fragment Analyzer (Agilent). miRNA libraries were constructed using the QIAseq miRNA UDI Library Kit (QIAGEN, 331502). Briefly, pre-adenylated 3’ adapters were ligated to the 3’ ends of miRNAs, followed by 5’ adapter ligation. The ligation products were reverse-transcribed into cDNA using an RT primer containing unique molecular indexes (UMIs) and purified with magnetic beads. The purified cDNA was amplified by PCR using primers with unique dual indexes (UDIs), followed by a second bead purification to yield ∼200 bp miRNA sequencing libraries. The libraries were qualified using the Qubit™ 4 fluorometer (Thermo Fisher Scientific) and assessed for size distribution using 5200 Fragment Analyzer (Agilent).

Proteins were extracted with urea containing lysis buffer (4 M urea, 50 mM EDTA, 1 mM PMSF and 5 mM DTT) through repeated sonication and liquid nitrogen freeze thaw cycles. Following reduction (DTT) and alkylation (IAA), the samples were processed via 10 kDa ultrafiltration according to the FASP workflow. Proteins were digested with 50 mM NH₄HCO₃ containing 20 ng/μL trypsin at 37 °C for 16 h, and the resulting peptides were desalted using C18 StageTips. Peptides were separated on an Easy nLC system on a C18 analytical column using a 65 min gradient and analyzed on Orbitrap Exploris 480 mass spectrometer in data dependent acquisition (DDA) mode with HCD fragmentation. Raw files were processed with PEAKS 8.5 software (Bioinformatics Solutions Inc., Canada). De novo sequencing was performed first, and peptide identifications with an ALC Score≥50 were retained. Then raw files were then searched against the Staphylococcus aureus protein database downloaded from Uniprot (taxonomy ID: 1280, download on May, 2022), to which 286 common contaminant proteins were appended. Searching parameters were set as follows: enzyme, trypsin; precursor and fragment ion mass tolerance was set to ±10 ppm and ±0.02 Da; variable modification, oxidation of methionine and deamidation of asparagine and glutamine; fixed modification, carbamidomethylation of cysteine. A cutoff of –10lgP ≥15 was applied to remove low-confidence peptide identification, and only proteins with –10lgP ≥20 were retained. Contaminant peptide and protein identifications were removed before downstream analyses. The numbers of the identified peptides and proteins were counted.

Based on the measured RNA content, the number of identified peptides, and the estimated total volume of membrane microspheres calculated assuming a diameter of 600nm), the concentrations of the RNA and peptides in membrane microspheres were estimated.

### 6. Morphological characterization and size distribution analysis of lipid bilayer membrane microspheres by cryo-transmission electron microscope

The sample was mixed with an equal volume of physiological saline, and 2 μL of the mixture was added onto a glow-discharged holey carbon grid (Quantifoil Cu R1.2/1.3, 300 mesh). After incubation for 3 minutes at room temperature, the grid was plunge-frozen in liquid ethane using a Vitrobot Mark IV cryo-electron microscopy sample preparation instrument (FEI/Thermo Fisher Scientific) under 95% humidity and 4℃, with the following blotting conditions: blot time = 6s, blot fore = 15, wait time = 3s, blot total = 1. The frozen grid was stored in liquid nitrogen until use. For imaging, the copper grids were transferred into the cryo-holder of a Talos F200C cryo-transmission electron microscope (FEI/Thermo Fisher Scientific) equipped with a Ceta 4k × 4k camera at the Center of Cryo-Electron Microscopy, Zhejiang University (Hangzhou, China). Lipid bilayer membrane structure was verified at high magnification by the characteristic “dark-light-dark” pattern in the peripheral region of particles. Low-magnification imaged were then taken for each sample, and particles larger than 80nm were classified and counted. Particles in which the peripheral lipid bilayer membrane accounted for more than 70% of the circumference were classified as lipid bilayer membrane microspheres, and their proportion relative to the total counted particles was calculated.

### 7. B16-F10 cell and culture

The murine melanoma cell line B16-F10 (catalog no. SCSP-5233) was obtained from the Cell Bank of the Committee for Type Culture Collection of the Chinese Academy of Sciences (Shanghai, China). Cells were cultured in RPMI 1640 medium (Gibco) supplemented with 10% fetal bovine serum (FBS) and maintained in a humidified incubator at 37 °C under 5% CO₂. All experiments were performed using cells in the logarithmic growth phase with viability greater than 95%.

### 8. B16-F10 cell suspension preparation and storage

B16-F10 cells in the logarithmic growth phase with viability greater than 95% were washed three times with sterile PBS, digested with trypsin, and the digestion was terminated with complete medium, followed by gentle pipetting to obtain a single-cell suspension. The suspension was centrifuged at 200 × g for 5 min, the supernatant was discarded, and the cell pellet was collected. An aliquot of the cells was subjected to acridine orange (AO)/propidium iodide (PI) dual-fluorescence staining combined with an automated cell counter to determine cell viability: 10 μL of the cell suspension was mixed with 10 μL of AO/PI working solution (final concentrations of both AO and PI: 5 μg/mL, prepared in sterile PBS) and incubated at room temperature in the dark for 3–5 min; 12 μL of the mixture was then loaded into the cell counter for counting. If the cell viability was greater than 90%, the cell suspension was adjusted to the desired concentration (e.g., 1 × 10⁶ cells/mL) with sterile deionized water according to the experimental requirements. The vials containing the adjusted cell suspension were sealed with parafilm, the storage box was completely wrapped in aluminum foil to protect the samples from light, and the samples were finally stored at 37 °C.

### 9. Quantification of viable B16-F10 cells and determination of time to CCD

After incubation at 37 °C, samples were taken at the indicated time points (immediately and at 20, 40, 60, 80, 100, 120, 140, 160, 180, 240, 300, 360, and 420 min). For each time point, 10 μL of the cell suspension was mixed with 10 μL of AO/PI working solution (final concentrations of both AO and PI: 5 μg/mL, prepared in sterile PBS) and incubated at room temperature in the dark for 3–5 min; 12 μL of the mixture was then loaded into an automated cell counter to determine the ratio of live to dead cells. In parallel, a clonogenic (single-cell colony formation) assay was performed to further evaluate cell survival and proliferation capacity: 500 μL of the cell suspension was added to 5 mL of RPMI 1640 medium containing 20% FBS and cultured in an incubator at 37 °C with 5% CO₂ for 20–30 days, with half of the medium replaced every 3–4 days. At the end of the culture period, the culture medium was removed, the cells were washed once with PBS, and fixed with 4% paraformaldehyde for 10–20 min. Colonies were stained with 0.5% crystal violet solution (prepared in methanol) at room temperature for 10–30 min, rinsed under running water to remove excess stain, air-dried, photographed, and counted. Cell viability was comprehensively evaluated by combining these two methods to confirm that the cells met the viability requirements for subsequent experiments.

## Results and Discussions

### The time course of complete cell death of *Staphylococcus aureus (S. aureus)* in pure liquid water

From February 2014 to July 2015, we observed complete cell death (CCD) of *S. aureus* placed in the sealed system of pure water. *S. aureus* exhibited robust growth on standard culture medium, producing colonies 1–2 mm in diameter that were milky-white and glossy, circular and convex, with smooth, non-spreading margins (**Figure 1a**). Cells were harvested at the logarithmic growth phase and subjected to three successive washes with physiological saline via centrifugation. The resulting bacterial pellet was resuspended in pure water and aliquoted into 2-ml sterile cryogenic vials at a volume of 1.8 ml per vial (**Figure 1b**). The vials were then sealed, protected from light (**Methods**), and maintained at room temperature. Each vial contained an average bacterial load of 2.55 × 10⁹ cells, with the bacterial suspension appearing as turbid (Method)

**Figure 1.**
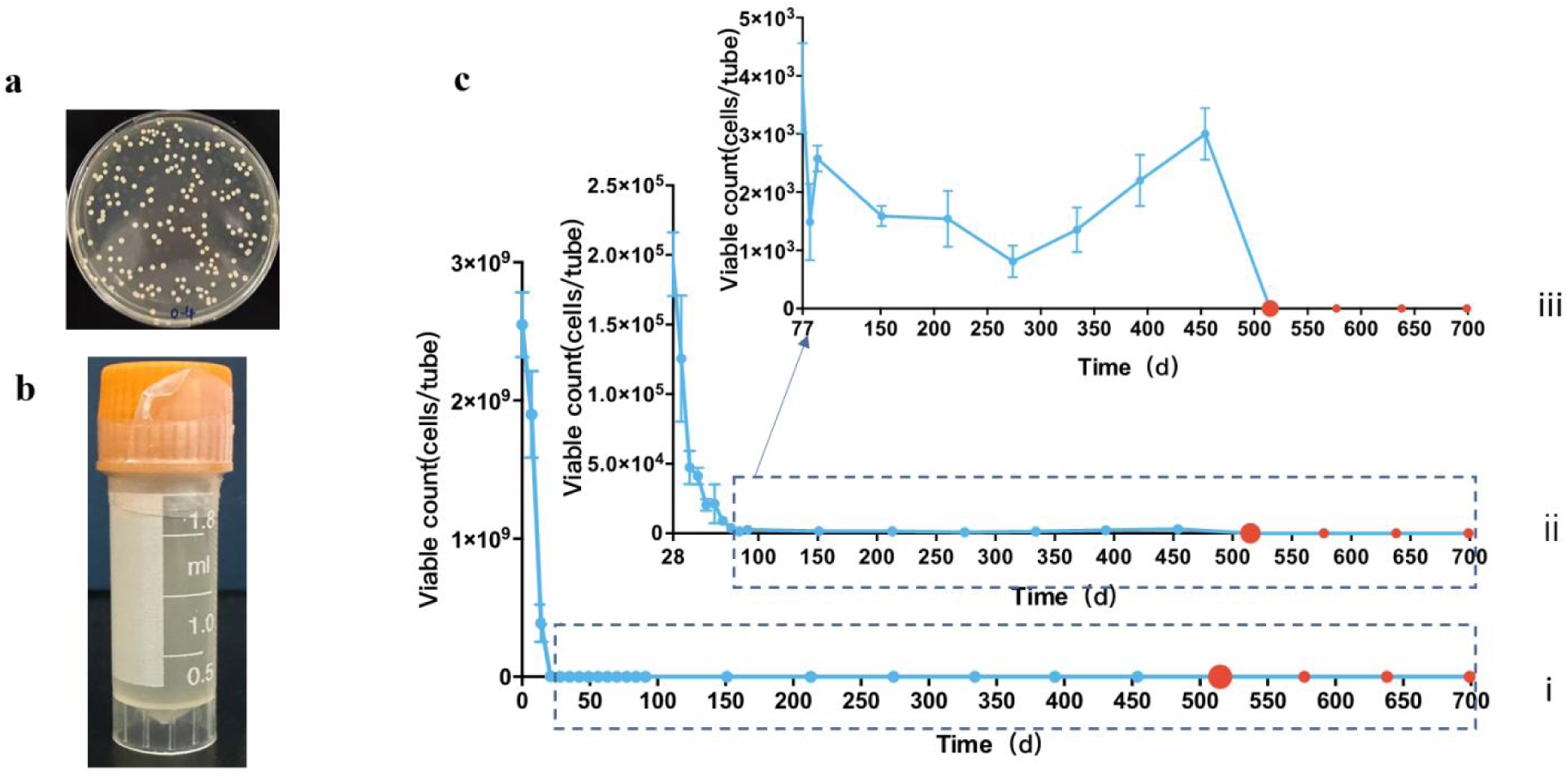
Survival dynamics of 2. 55×10^9^ *S. aureus* cells in sealed vials of pure water under room temperature. **a,** Colony morphology of *S. aureus* on LB agar plates. Colonies were circular, smooth, and white. Culture dish diameter: 90 mm. **b,** *S. aureus* suspension storage vials. Commercial 2-ml cryogenic vials, each loaded with 1.8 ml of bacterial suspension; screw caps were tightened and the vials were sealed with adhesive sealing tape. **c,** Survival curves depicting viable bacterial counts in sealed vials over time. **(i)** presents the original survival curve spanning days 0–699, **(ii)** shows a magnified view of the survival curve from days 29–699, **(iii)** presents a further magnified view from days 77–699. Viable bacterial counts reached zero on day 515 (first red dot), with subsequent assays at equal intervals on days 577, 638, and 699 (red dots) consistently returning zero, respectively. The x-axis denotes time (days); the y-axis denotes the number of viable *S. aureus* cells

We define complete cell death (CCD) of *S. aureus* as the state in which viable cell counts decline to zero after static placement in the sealed vials (Methods). To determine the time to CCD for *S. aureus*, we randomly sampled 3 vials at irregular time intervals and enumerated the viable bacterial counts therein (**Methods**). The findings revealed that, from an initial inoculum of 2.55×10^9^ cells per vial, the viable count declined to 1.93×10^5^ ± 2.29×10^4^ by day 28 (**Figure 1c(i)**), dropped to 3.79×10^3^ ± 7.70×10^2^ by day 77 (**Figure 1c(ii)**), and subsequently decreased gradually with minor fluctuations between day 77-454, when it reached zero on day 515 (**Figure 1c(iii)**). Thereafter, three consecutive assays performed at a 61-day interval consistently scored zero viable cells. Accordingly, day 515 was designated as the definitive time point of CCD of *S. aureus* in the sealed system that were kept at room temperature (**Figure 1c(iii)**).

Both bacterial density and ambient temperature within the sealed vials affected the time to CCD of *S. aureus*. At room temperature, an inoculum of 1.91×10^8^ cells per vial required 280 days to reach CCD (**Extended Data Figure 1**)—235 days shorter than that observed for 2.55×10^9^ cells per vial. Thus, higher bacterial densities necessitated proportionally longer time intervals to achieve CCD. Under constant 37°C, the inoculum of 3.02×10^8^ cells per vial exhibited a markedly abbreviated time to CCD, shortening to 21 days (**Extended Data Figure 2**). By contrast, at 2.55×10^8^ cells per vial maintained under constant 25°C, 9.08×10^3^ viable cells still persisted at day 1456.

Across the bacterial densities and ambient temperature conditions examined, we consistently observed that the majority of bacteria within the sealed vials succumbed rapidly. A small subpopulation sustained viability, probably by deriving substrates from the degradation of earlier deceased cells, thereby sustaining metabolic activity for a limited duration; yet, they were ultimately destined to forfeit the capacity to uphold their own stability in water upon metabolic exhaustion.

And so we asked: what persists in the CCD suspension—kept in a sealed system and left to stand over a long duration of time at room temperature? To fathom this, we established a long-term continuous observation cohort – 6.38 × 10⁹ cells per vial, from December 2024 onward—for the inquiries that follow.

## A class of lipid bilayer membrane microspheres in the post-CCD suspension

A suspension sample from the day-3423 vial of the room-temperature long-term observation group was examined by cryo-electron microscopy (cryo-EM). We found that the observed particles were almost exclusively a class of lipid bilayer membrane microspheres (abbreviated as “membrane microspheres”), with diameters ranging from 80-600nm (**Figure 2**). Of the 2,281 particles visible in the field of view, 2,133 were lipid bilayer membrane microspheres (93.51% of the total), and 148 were non-lipid bilayer membrane microsphere structures (6.49%; **Extended Data Fig. 3**). Cryo-EM examination of the original suspension from the day 4,248 vial similarly revealed almost exclusively membrane microspheres; of 1,064 particles counted in the field of view, microspheres accounted for 95.11%.

**Figure. 2.**
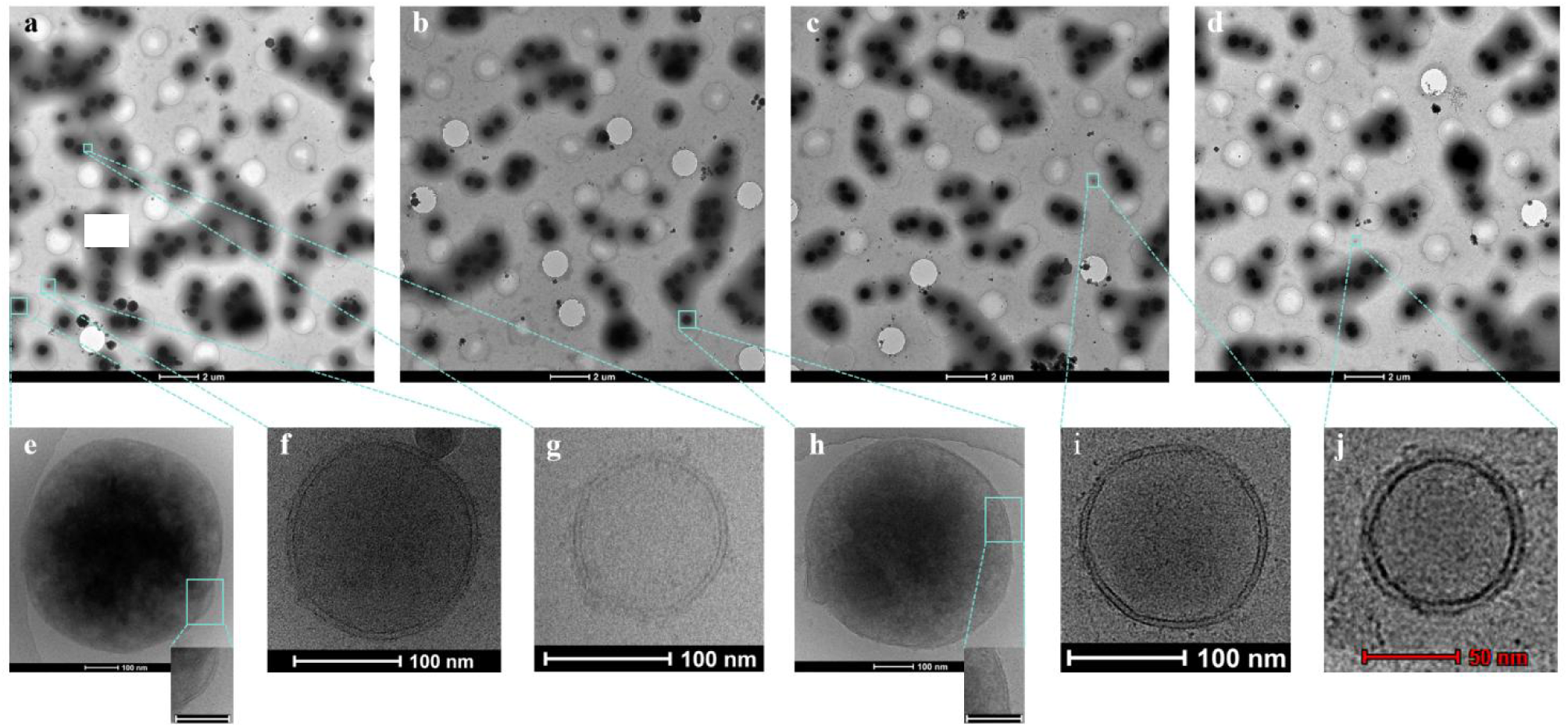
Cryo-TEM morphology of lipid-bilayer membrane microspheres in the CCD suspension of the long-term observation group at day 3423, room temperature. **a–d**, Abundant microspheres of varying sizes were visible in the fields of view; no dead *S. aureus* remnant structures were observed. Scale bar, 2 µm. **e–j**, Magnified views revealing that these microspheres exhibited typical lipid bilayer membrane structures. Scale bars: 100 nm for e–i; 50 nm for j.

Cryo-EM examination of original suspension samples spanning a decade demonstrates that the CCD solution within the sealed system — once filled with bacterial structures—has transformed into a suspension almost exclusively comprising of membrane microspheres. What, then, constitutes the self-purification process by which these membrane microspheres emerge within the CCD suspension?

Accordingly, we conducted self-purification experiments of membrane microspheres under constant 37°C—a condition that enables the most rapid attainment of CCD—using two inoculum densities (3.02 × 10⁸ and 4.10 × 10⁸ cells per vial). Cryo-EM examination of these two CCD suspensions revealed that, in one group, the proportion of lipid bilayer membrane microspheres increased from 25.58% (day 182) to 50.62% (day 983); in the other group, it rose from 48.92% (day 324) to 72.76% (day 1,125). **Figure 3** illustrates the progressive degradation of bacterial structures within the CCD suspension along the time axis from day 182 to day 1,125.

**Figure. 3.**
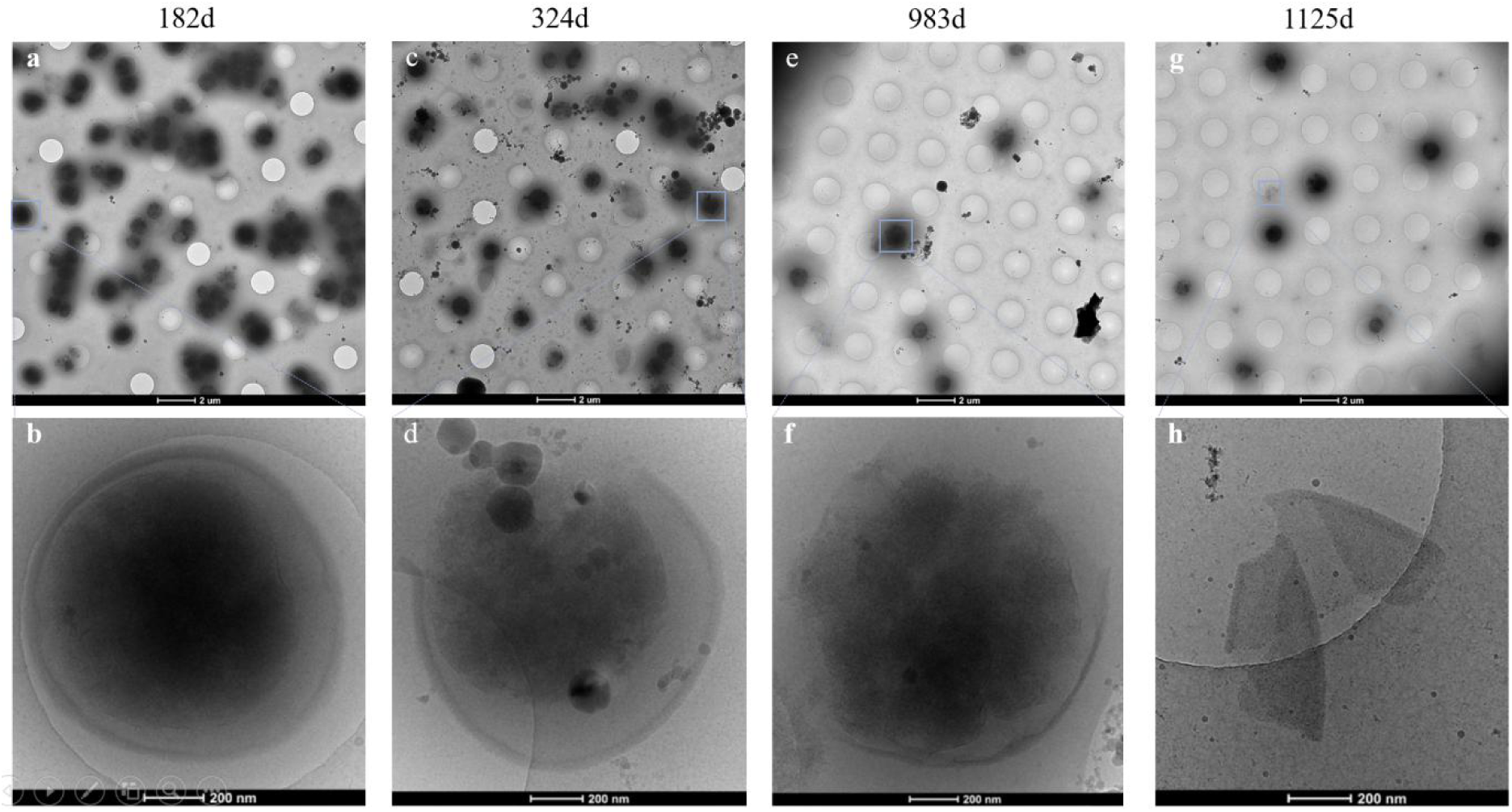
Cryo-EM morphology of the degrading *S. aureus* cell remnants in sealed system at a constant 37 °C. **a,** Day 182: the field was crowded with dead *S. aureus* remnants. Scale bar, 2 µm. **b,** Magnified view at day 182: some remnants still held an intact cell wall. Scale bar, 200 nm. **c,** Day 324: the remnants remained visible, yet fewer. Scale bar, 2 µm. **d,** Magnified view at day 324: some cell walls were now partially degraded. Scale bar, 200 nm. **e,** Day 983: only a small number of remnants remained in view. Scale bar, 2 µm. **f,** Magnified view at day 983: a few remnants retained incompletely degraded cell walls. Scale bar, 200 nm. **g,** Day 1125: no cell wall-bearing remnants could be found in the field. Scale bar, 2 µm. **h,** Magnified view at day 1125: only degraded, detached fragments of cell walls remained—evidence of what has already vanished. Scale bar, 200 nm.

These observations tell of two fates in the pure water in the sealed system after CCD: upon CCD, the dead bacterial structures are progressively degraded with increasing time – owing to their inherent instability; by contrast, the membrane microspheres arising from the CCD can maintain long-term stability, achieving self-purification once the cellular structures have been completely degraded.

In other words: given sufficiently long storage time, the CCD suspension in the sealed system will in the end hold almost nothing else, but only such membrane microspheres.

The phenomenon of reversion to such lipid bilayer membrane microspheres upon CCD in prokaryotic cells likewise been validated in eukaryotic (**Extended data Figures 4-5**).

### Dynamic variation in the abundance and size of membrane microspheres in the CCD suspension

After more than 2,000 days of undisturbed storage in darkness at ambient temperature, we serendipitously observed that the cell suspensions within the sealed systems of the long-term observation group—originally opaque and quicksand-like in appearance—had become limpid and transparent. Accordingly, between days 2310 and 4059, we randomly sampled specimens at eight irregularly spaced time points, and employed nano-flow cytometry to quantify the abundance and size distribution of suspended particles. The data demonstrated that the total number of particles in the sealed systems varied in a fluctuating manner over the observation period (**Figure. 4a, b**). The total particle counts per vial were: 1.11× 10^9^ on day 2310, 1.68× 10^9^ on day 2375, 4.78× 10^9^ on day 2408, 4.56× 10^9^ on day 2424, 7.34×10^8^ on day 2798, 8.30×10^8^ on day 3215, 8.75×10^8^ on day 3423, and 2.34× 10^9^ on day 4059. For analysis, we divided the total particle population of each sealed system—by measured size alone, an arbitrary partition—into three size ranges: large (φ > 590 nm), medium (300 nm < φ ≤ 590 nm), and small (80 nm < φ ≤ 300 nm), φ being the diameter. Along the temporal axis, the particle counts corresponding to these three size ranges exhibited pronounced dynamic fluctuations—increasing or decreasing substantially across time points (**Figure. 4b**). Proportion analysis of the differently sized particles in the sealed system at specific time points revealed that, across the eight observation time points, small particles accounted for 20.51%–77.80%, medium particles for 20.77%–79.04%, and large particles for a relatively minor fraction (0.04%–3.02%) of the total, respectively (**Figure. 4c**). Thus, the particle population in the sealed system was dominated by small– and medium-sized particles, whose relative proportions fluctuated markedly over time—indicating that the size distribution of particles within the sealed system exhibits significant temporal dynamics and that interconversion among size classes may occur.

**Figure. 4.**
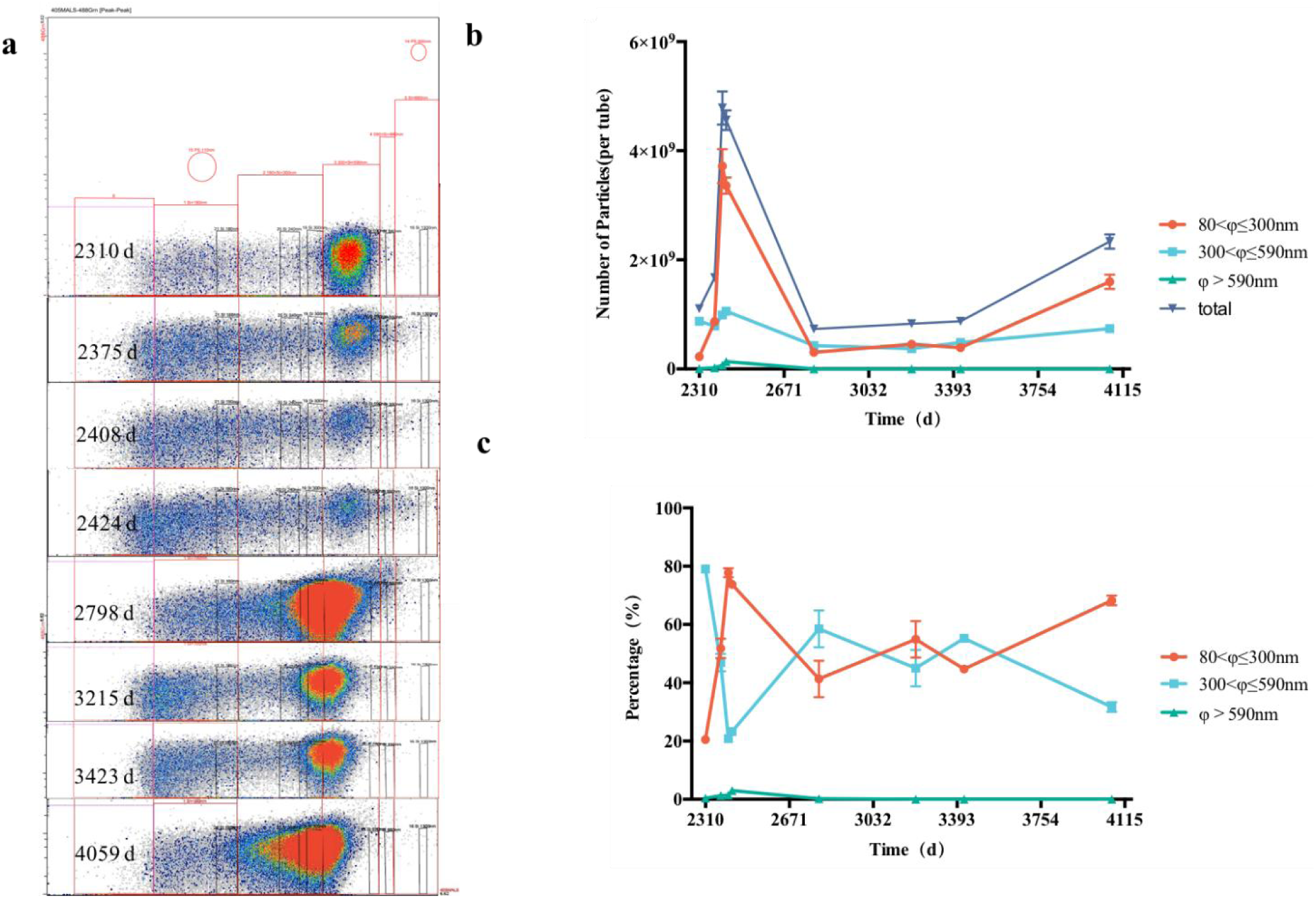
Dynamic changes in particle abundance and size in the CCD suspensions of *S. aureus* from the long-term observation group at room temperature. **a,** Particle size distributions of the *S. aureus* CCD suspension in the sealed system, measured by the Apogee ultrasensitive nano-flow cytometer. From top to bottom, stacked plots show the particle distributions at days 2310, 2375, 2408, 2424, 2797, 3215, 3423, and 4059, respectively. The stacked plots visualize the relative abundance of particle events across size fractions, with colors from gray, blue, green, yellow to red representing increasing event abundance. The y-axis donates 488-Grn (reflecting particle purity) and the x-axis donates MALS middle-angle light scattering (reflecting particle size). From left to right, six rectangular gates (red rectangles) define the analysis regions: the leftmost gate excludes instrument background noise, while the remaining five gates are drawn according to the positions of ApogeeMix standard microspheres in the side-scatter channel, corresponding to the 80– 180 nm, 180–300 nm, 300–590 nm, 590–880 nm, and >880 nm size ranges, respectively. Particle abundance was high in the 80–180 nm, 180–300 nm, and 300–590 nm regions, with marked temporal variation in abundance within each region at different time points. **b,** Temporal variation curves of the total particle count and of particle counts within the large (φ > 590 nm), medium (300 nm < φ ≤ 590 nm), and small (80 nm < φ ≤ 300 nm) size ranges (φ denotes diameter) in the sealed system across the eight time points. The x-axis donates time (days) and the y-axis donates absolute particle counts (particles per tube). Data are presented as mean ± SD (n = 3). The total particle counts and the counts of medium– and small-sized particles exhibited pronounced dynamic increases and decreases across time points; the medium particles fluctuated between 20.77% and 79.04% and the small particles between 20.51% and 77.80%, whereas the large particles (>590 nm) fluctuated within a narrow range of 0.04%–3.02%. **c,** Time-series line plot of the percentage of particles within the large, medium, and small size ranges in the sealed system. Particles smaller than 590 nm—i.e., the medium and small particles—constituted the dominant particle population in the sealed system. Across the eight time points, the percentage of medium particles was 79.04%, 46.98%, 20.77%, 23.25%, 58.42%, 45.18%, 55.25%, and 31.62%, while that of small particles was 20.51%, 51.72%, 77.80%, 73.73%, 41.38%, 54.75%, 44.69%, and 68.34%; the two curves crossed at multiple time points, indicating alternation of dominance between the two-size population of particles. The percentage of large particles, bounded by day 2424, rose from 0.45% to 3.02% in the earlier phase and declined to 0.04% thereafter.

In view of the strictly sealed nature of our experimental system—wherein no external matter or energy is introduced, apart from passive thermal tracking of ambient temperature (From December 2014 to June 2026, monthly mean temperatures in Chongqing, China, ranged from a minimum of 7.5 °C to a maximum of 35.3 °C). Thus, we have grounds to posit that the energy that drives the dynamic fluctuations in the size and abundance of the membrane microspheres in the CCD suspension could originate solely from the thermal energy of the motion of water molecules driven by the ambient temperature changes.

Similar dynamic fluctuations in the size and abundance of membrane microspheres were observed in the post-CCD suspensions analyzed in short-term supplementary experiments (room temperature, **Extended Figure 6**; 37 °C, **Extended Figure 7**) via nanoflow cytometry. Furthermore, the proportion of the small and medium-sized particles relative to the total particle count in the sealed systems rose almost from zero at the outset, increasing progressively throughout the CCD process (**Extended Figure 6c, Extended Figure 7c**), indicating that the vast majority of the particles in these two size ranges were generated *de novo* during the CCD process.

### RNA and protein composition of membrane microspheres in the CCD suspension

For the long-term continuous observation group, we separated the membrane microspheres from the extra-microsphere solution of the 4,248-day sealed system and subjected both fractions to RNA and protein analysis. Both the membrane microspheres and the extra-microsphere solution were found to contain trace amounts of RNA. Capillary electrophoresis revealed that the RNAs from the two sources both exhibited single-peak distributions, peaking at 87 nt and 77 nt, respectively. RNA libraries constructed under comparable conditions further showed that membrane microsphere-derived RNA displayed a broader size distribution and higher abundance than that from the extra-microsphere solution. Per unit volume, the RNA concentration within the membrane microspheres was estimated to be 36.8-fold higher than that of the extra-microsphere solution. Mass spectrometric analysis identified 1,054 proteins in the membrane microsphere samples and 567 in the extra-microsphere solution samples; based on the number of detected peptides per unit volume, the protein concentration of the membrane microspheres was estimated to be 146.32-fold higher. We posit that the biomolecules detected in the membrane microspheres group represent, for the most part, remnants of products synthesized by *S. aureus* in accordance with the central dogma of molecular biology. Equally, we foresee that these membrane microspheres may also harbor nucleotide chains and peptide chains that do not conform to the central dogma.

Jim Al-Khalili articulates that life is able to sustain its living state; so, the minimum requirement we will demand for our bottom-up protocells to earn classification as living is that they must be capable of sustaining themselves in turbulent thermal-dynamic seas^1^. That is to say, the capacity to withstand the destructive forces of aqueous thermodynamic fluctuations is the necessity for a primordial life form to be deemed alive.

Seawater contains abundant bacterial lipid bilayer vesicles, at concentrations reaching ∼10⁵ ml scale, and with marked structural stability^2^. We can foresee that simpler animate beings that predated the dawn of cellular life may yet endure, nonetheless its forms unfolded by classical Darwinian evolution across the ages of a changing Earth.

## Conclusion

By classical thermodynamic necessity, within the sealed systems of this study, the CCD suspension under aqueous thermodynamic storms is destined to resolve into a uniform solution of organic molecules. Yet over more than a decade of continuous observation, we discovered a class of “living” structures in the CCD suspension: microspheres bounded by lipid bilayer membranes, containing RNA and protein molecules, capable of harnessing thermal energy to maintain their long-term sustainability and stability in liquid water, and undergoing dynamic fluctuations in size and in abundance. Here, we designate this category of the lipid bilayer membrane microspheres as “Meta Bodies” (MB, ⊙). Meta bodies, perhaps, is a form of animate being that is simpler than the cell itself.

## End notes

### Authors’ contributions

Z.B. Wang conceived the study, acquired funding, provided resources, led the project, and provided critical review, discussion and revision of the manuscript. L.Q. Fang designed the experiments, organized and supervised the research, and wrote the original draft. Y.T. Wang developed and optimized the methods, conducted the investigation, curated and visualized the data, performed formal analysis, and prepared the samples. H. Qiao designed the morphological analysis methods and performed cryo-EM observation and RNA measurement. J. Ma performed cryo-EM morphological identification, proportion data analysis, and protein measurement. D.G. Yu and Y.Z. Jiang conducted the investigation and curated the data. J. Bai contributed to the conceptualization of the study, provided organizational support to the research team throughout the study, and critically reviewed and discussed the manuscript. Q.Y Xia contributed to the formal analysis, critically review and revision of the manuscript. H. Chen, D. Song and W.J. Song participated in the critical discussion of the results and revision of the manuscript. All the authors agreed to the manuscript’s publication

## Acknowledgments

We thank Professor Rosie Xing for thorough checking up of the original data, critically reviewing and revising the manuscript and for assistance with English translation; Dr. Shizhu Li for assistance with statistical analysis of the nanoflow cytometry data; Dr. Wen Xu for the constructive discussion of the study; and Lingyun Wu at the Center of Cryo-Electron Microscopy (CCEM), Zhejiang University, for technical assistance with cryo-EM.

## Availability of data and materials

The data that support the findings of this study are available from the corresponding authors upon reasonable.

## Supplemental information

Extended Figure 1-7

