## Extended Figure 1-7 for "Reversion to the Meta Body Form of Life after Complete Cell Death in Pure Liquid Water"

### Extended Data

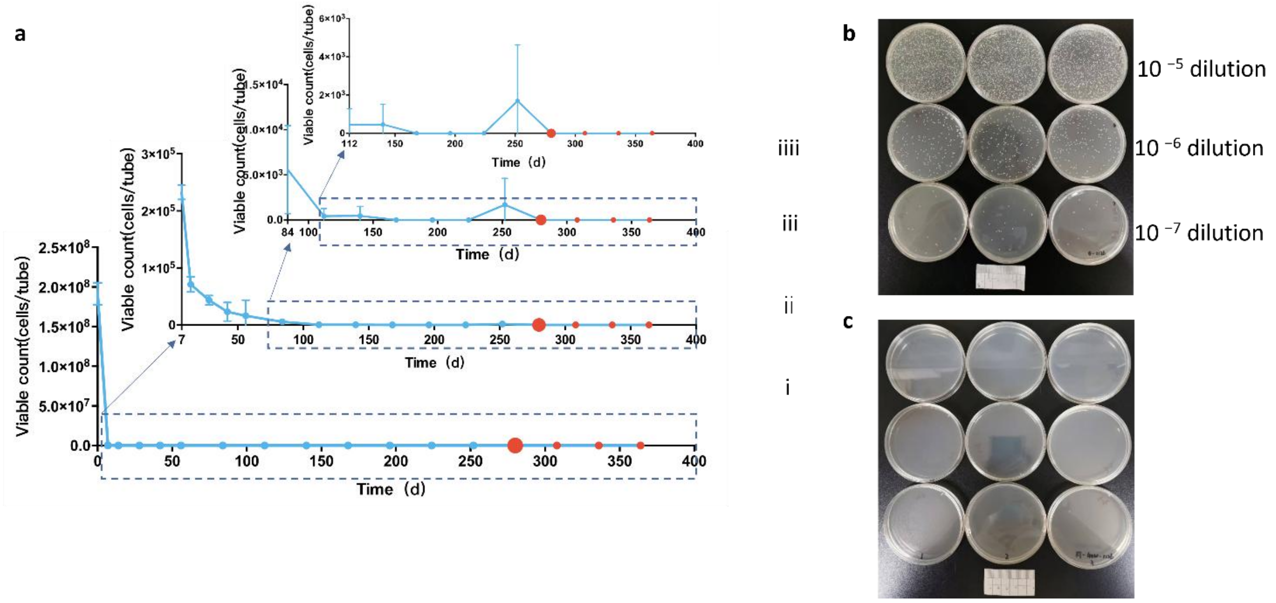


**Extended Data Fig. 1 | Changes in 1.91 x $10^8$ viable *S. aureus* cell counts in sealed vials of pure water at room temperature.** **a,** Survival curves showing viable bacterial counts over time in sealed vials under room-temperature conditions. **(i)** Full-range curve spanning days 0–364**. (ii)** Magnified view of the survival curve from days 7–364. (iii) Magnified view of the survival curve from days 84–364. **(iv)** Magnified view of the survival curve from days 121–364. Viable counts reached zero on day 280 (first red dot), with three consecutive assays at equal intervals on days 308, 336, and 364 (red dots) consistently returning zero, respectively. The x-axis denotes time (days); the y-axis denotes viable *S. aureus* counts per vial (cells/vial). **b,** Colony morphology of *S. aureus* on LB agar plates at the initial aliquoting time point following serial dilution ($10^{-5}$, $10^{-6}$, and $10^{-7}$ from top to bottom; n = 3). 90-mm culture dishes. **c,** Colony morphology of *S. aureus* on LB agar plates on day 280 by full plating of the sample (n = 3). Colony formation was not detected. 90-mm culture dishes.

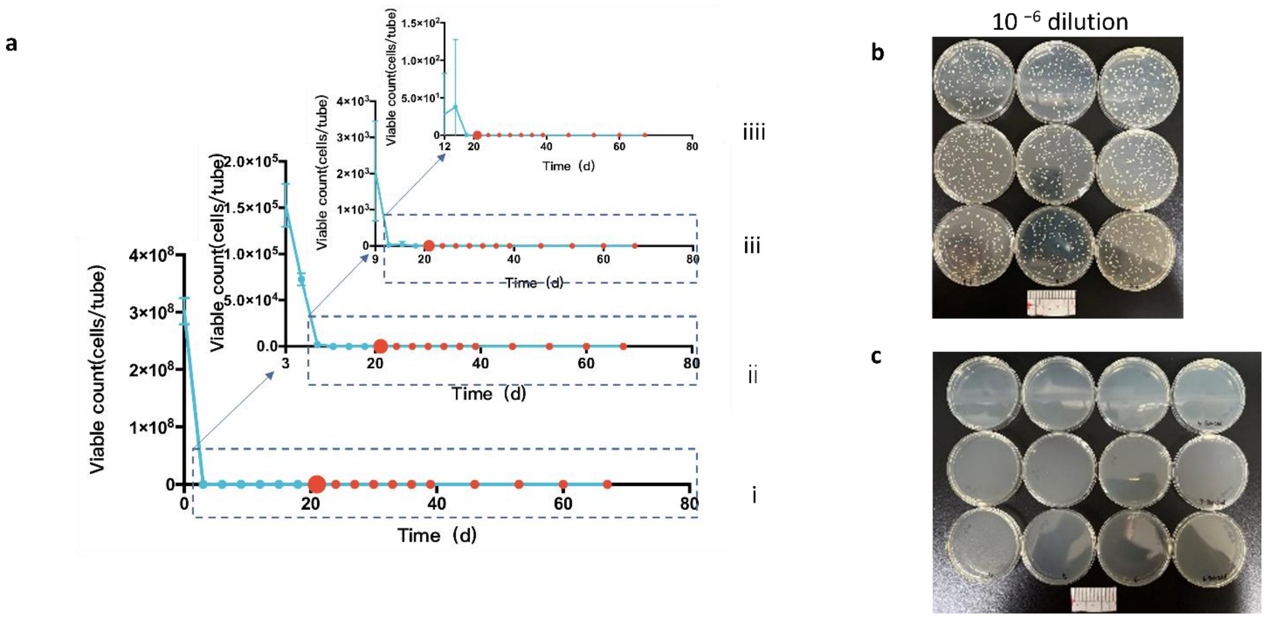


**Extended Data Fig. 2 | Changes in 3.02 x $10^8$ viable *S. aureus* cell counts in sealed vials of pure water at 37°C. a,** Survival curves showing viable bacterial counts over time in sealed vials under 37°C. **(i)** Full-range curve spanning days 0–67. **(ii)** Magnified view of the survival curve from days 3–67. **(iii)** Magnified view of the survival curve from days 9–67. **(iv)** Magnified view of the survival curve from days 12–67. Viable counts reached zero on day 21 (the first red dot). Viable bacteria remained undetectable at all subsequent time points—24, 27, 30, 33, 36, 39, 46, 53, 60, and 67 days (red dots). The x-axis denotes time (days); the y-axis denotes viable *S. aureus* counts per vial (cells/vial). **b,** Colony morphology of *S. aureus* on LB agar plates at the initial aliquoting time point following $10^6$ dilution（n = 3). 90-mm culture dishes. **c,** Colony morphology of *S. aureus* on LB agar plates on day 21 by full plating of the sample (n = 3)). Colony formation was not detected. 90-mm culture dishes.

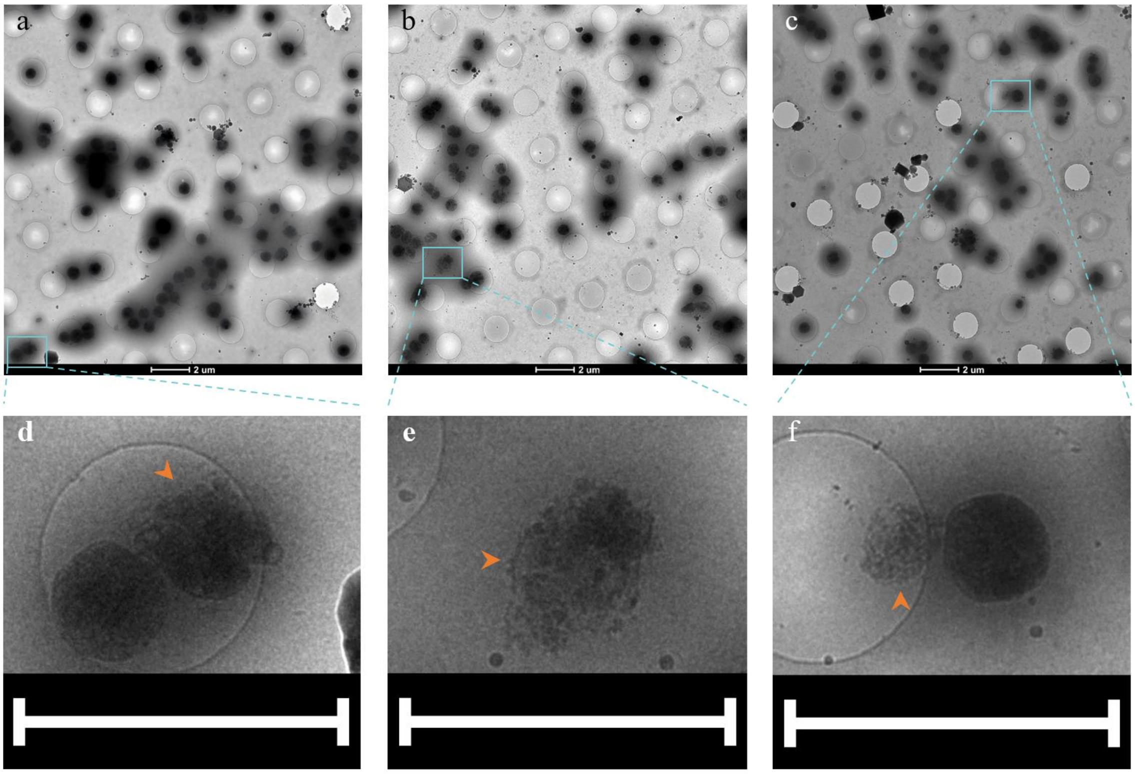


**Extended Data Fig. 3 | Cryo-transmission electron microscopy (cryo-TEM) analysis of non-lipid-bilayer-membrane microspheres in the day-3423 sample of the long-term observation group. a–c,** A trivial number of particles other than lipid bilayer membrane microspheres were visible in the day 3423 sample. Scale bars, 2μm. **d–f,** Magnified views of these particles reveal two morphological classes (orange arrows): **d,** particles with severely deficient lipid bilayer membranes, and **e–f** aggregated particles devoid of lipid bilayer membranes. Scale bars, 2μm.

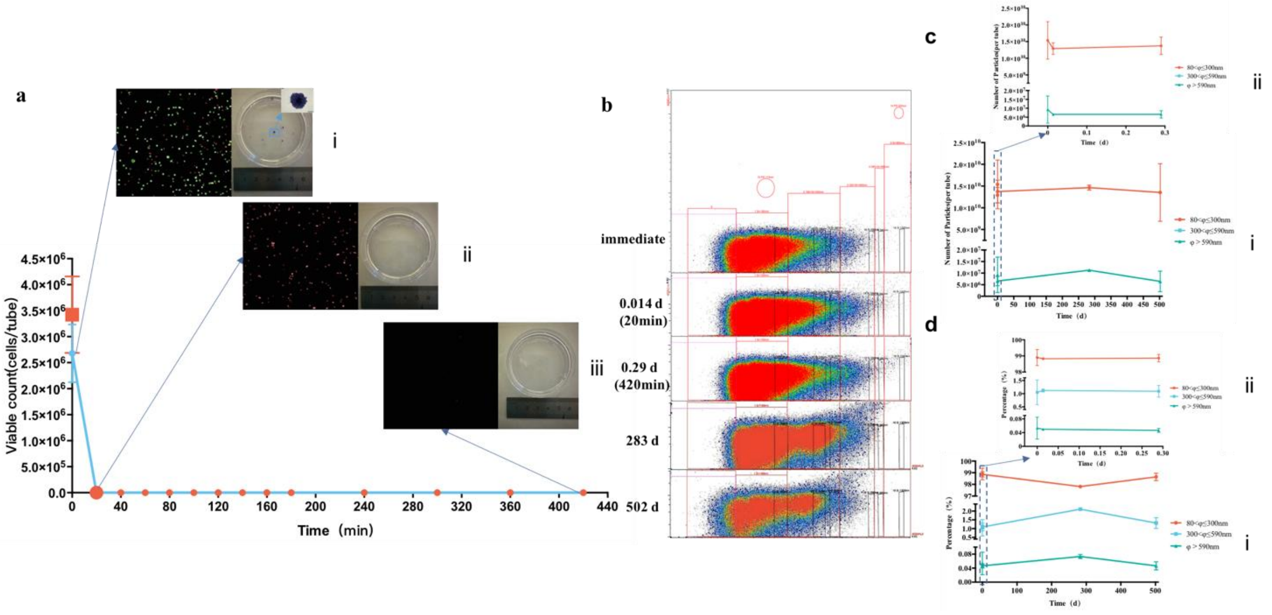


**Extended Data Fig. 4 | Time to complete cell death (CCD) and the dynamic changes in particle count and size of $10^6$-scale B16-F10 cells in pure water within sealed vials at 37 °C. a, T**emporal changes in the viability and clonogenic capacity of B16-F10 cells in the sealed vials. The curve shows the number of viable cells within the sealed vials over time under 37 °C conditions. Each vial initially contained $3.43 \times 10^6$ cells (red squares); the viable count was $2.68 \times 10^6$ immediately after suspension in pure water, dropped to zero at 20 min (first red dot), and remained zero at all subsequent time points of 40, 60, 80, 100, 120, 140, 160, 180, 240, 300, 360, and 420 min (red dots). **(i–iii)** Representative AO/PI fluorescence staining images (left) and corresponding cell clonogenic assay photographs (right) collected at the immediate time point, the CCD time point (20 min), and the end of the survival curve (420 min), respectively. In AO/PI staining, live cells appeared green (AO$^+$) and dead cells red (PI$^+$). In (i), the macroscopic view shows cell colonies obtained by $2 \times 10^5$-fold dilution of the immediate time-point sample (whole-well staining image, 60 mm culture dishes); the inset in the upper right corner is a bright-field micrograph of a single colony. AO/PI staining shows that the live-cell signal dropped sharply to zero at 20 min, while the dead-cell signal progressively diminished with prolonged storage; concomitantly, colony-forming capacity was completely lost at 20 min. 60 mm culture dish. **b,** Particle size distribution of B16-F10 suspensions measured by the Apogee ultrasensitive nano-flow cytometer. Stacked density plots from top to bottom show the particle distributions at the immediate time point and at 0.014 d (20 min, CCD), 0.29 d (420 min), 283 d, and 502 d, respectively. The stacked plots visualize the relative abundance of particle events across size gates, with colors from gray, blue, green, and yellow to red representing increasing event abundance. The x-axis donates the MALS multi-angle scatter intensity (reflecting particle size) and the y-axis donates the 488-Grn fluorescence intensity (reflecting particle purity). Particles were concentrated mainly in the 80–300 nm regions, with a slight shift to the left or right over time. **c,** The course of total particle counts and absolute counts within the three size ranges across the above time points. Each point is the arithmetic mean of three independent tubes, measured once per tube at each time point; counts are plotted on a logarithmic y-axis as $10^6$ particles per tube. Tick labels report the actual elapsed run time, with date values rounded for display. **d,** Time-series line plot of the percentage of particles in each size fraction at the indicated time points. The proportion of particles in each size interval fluctuated over time, with the direction of change inconsistent across fractions (i.e., not uniformly increasing or decreasing).

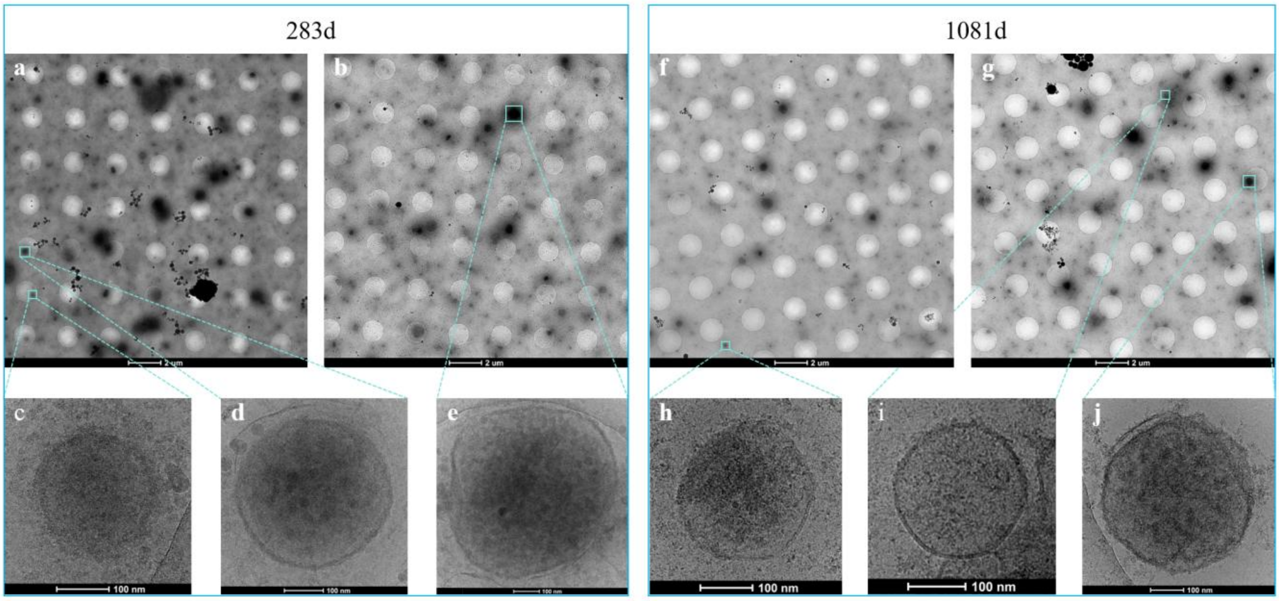


**Extended Data Fig. 5 | Cryo-EM morphology of lipid bilayer membrane microspheres in B16-F10 CCD suspensions within the sealed vial system at 37 °C. a–b**, Representative fields of the day-283 sample of the B16-F10 CCD suspension, showing numerous microsphere structures of various sizes. Scale bar, 2 μm. **c–e**, Magnified views of microspheres of different sizes, all exhibiting a lipid bilayer membrane structure, with diameters mainly distributed in the range of 80–600 nm. Scale bar, 100 nm. **f–g**, Representative fields of the day-1081 sample of the B16-F10 CCD suspension, showing numerous microsphere structures of various sizes. Scale bar, 2 μm. **h–j**, Magnified views of microspheres of different sizes, all exhibiting a lipid bilayer membrane structure, with diameters mainly distributed in the range of 80–600 nm. Scale bar, 100 nm.

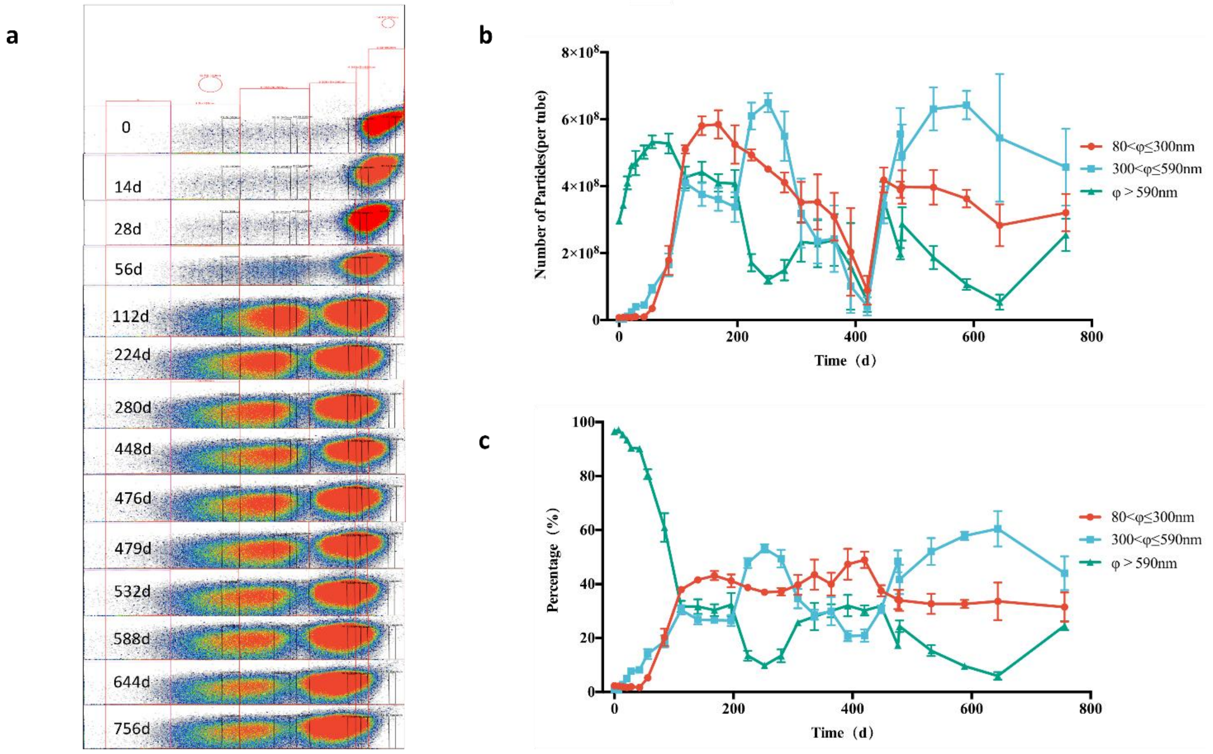


**Extended Data Fig. 6 | Dynamic changes in particle count and size of CCD suspensions of 1.91 × $10^8$ *Staphylococcus aureus* in sealed vials at room temperature. a** Particle size distribution of the *S. aureus* suspension measured by the Apogee ultrasensitive nano-flow cytometer. Stacked density plots from top to bottom show the particle distributions on days 0, 14, 28, 56, 112, 224, 280 (CCD time point), 448, 476, 479, 532, 588, 644, and 756, respectively. The stacked plots visualize the relative abundance of particle events across size gates, with colors from gray, blue, green, and yellow to red representing increasing event abundance. The x-axis donates the MALS multi-angle scatter intensity (reflecting particle size) and the y-axis donates the 488-Grn fluorescence intensity (reflecting particle purity). Marked changes occurred in the 80–300, and 300–590 nm fractions, where particle abundance was high. Initially, the particle population was concentrated at the large-size end (rightmost region); with prolonged storage, the population first shifted significantly to the left and then partially rebounded to the right, indicating that particle size changes were not unidirectional but underwent a non-monotonic, bidirectional dynamic remodeling. **b,** Temporal variation of particle counts within the large (φ > 590 nm), medium (300 nm < φ ≤ 590 nm), and small (80 nm < φ ≤ 300 nm) size ranges (φ denotes diameter) in the sealed vials at the above time points. The large, medium, and small particles fluctuated substantially in abundance—rising and falling markedly over time. Each data point represents the arithmetic mean of the averages of three independent tubes, with each tube's average calculated from three technical measurements. Counts are shown on a logarithmic y-axis as $10^6$ particles per tube, and the x-axis is proportional to elapsed time (days). **c,** Time-series line plot of the percentage of particles within the large (φ > 590 nm), medium (300 nm < φ ≤ 590 nm), and small (80 nm < φ ≤ 300 nm) size ranges in the sealed vials. The proportion of large particles (φ > 590 nm) declined sharply from 96.61% at the starting point to 13.43% at day 280 (the CCD time point). Correspondingly, the proportions of small particles (80 nm < φ ≤ 300 nm) and medium particles (300 nm < φ ≤ 590 nm) rose from 2.30% and 1.09% at the starting point to 37.17% and 49.39% at day 280, respectively. After a certain storage period, the percentages of large, medium, and small particles in the sealed vials fluctuated markedly over time.

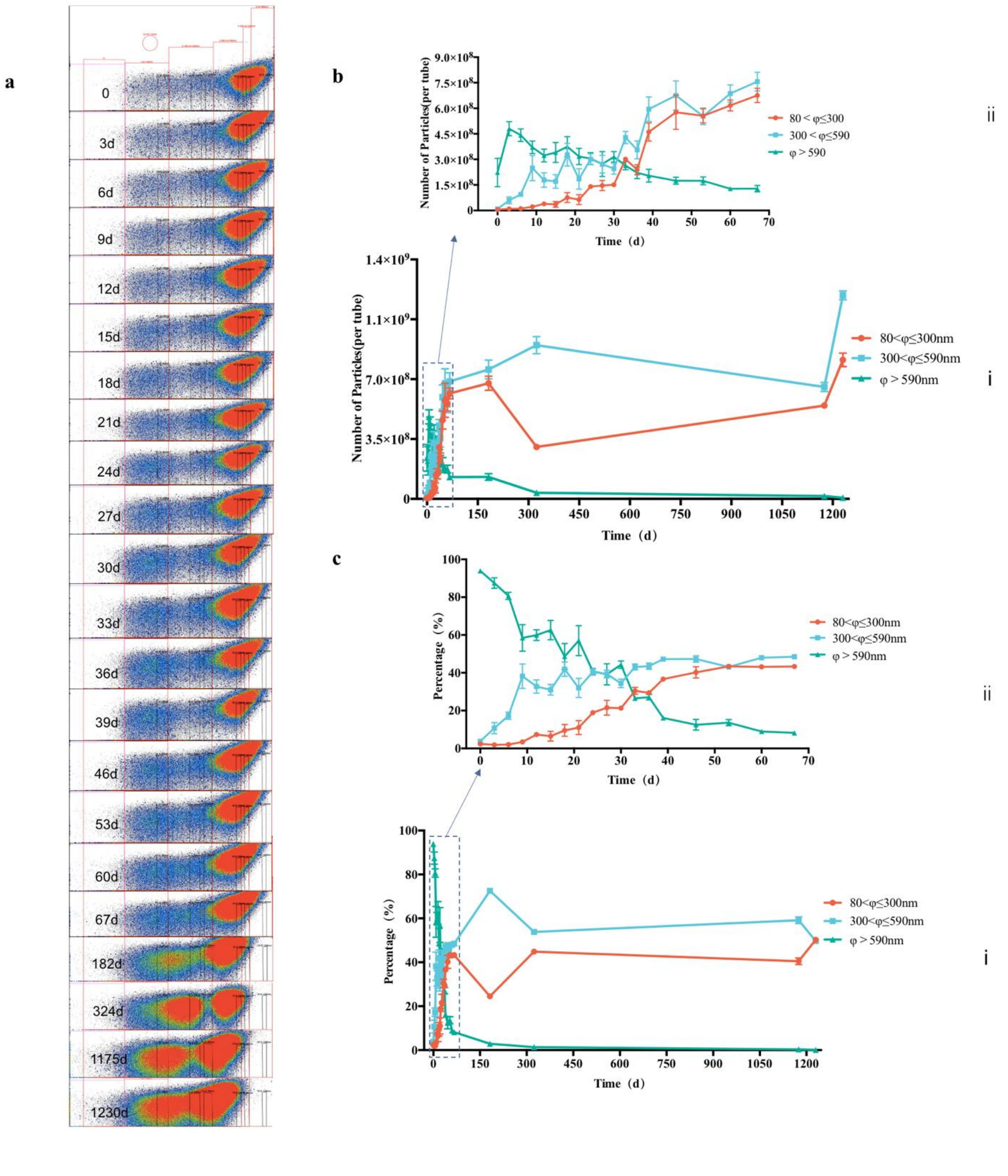


**Extended Data Fig. 7 | Dynamic changes in particle number and size of CCD suspensions of 3.02 × 10⁸ *Staphylococcus aureus* in sealed vials at 37 °C. a,** Particle size distribution of the S. aureus suspension measured by the Apogee ultrasensitive nano-flow cytometer. Stacked density plots from top to bottom show the particle distributions on days 0, 3, 6, 9, 12, 15, 18, 21 (CCD time point), 24, 27, 30, 33, 36, 39, 46, 53, 60, 67, 182, 324, 1175, and 1230, respectively. The stacked plots visualize the relative abundance of particle events across size gates, with colors from gray, blue, green, and yellow to red representing increasing event abundance. The x-axis donates the MALS multi-angle scatter intensity (reflecting particle size) and the y-axis donates the 488-Grn fluorescence intensity (reflecting particle purity). As shown in the figure, particle abundance exhibited a pronounced overall leftward shift (toward smaller sizes) with increasing storage time, with marked changes occurring in the 80–300 nm and 300–590 nm fractions. **b,** Temporal variation of total particle counts and absolute particle counts within the three size fractions at the above time points. Technical measurements within each tube were first averaged. (i) Each data point represents the arithmetic mean of the averaged values from three tubes at each sampling time point from day 0 to day 67, and the value of a single tube at days 182, 324, 1175, and 1230. Each tube was measured three times. Counts are shown on a logarithmic y-axis as $10^6$ particles per tube. (ii) An expanded view of days 0–67 using a continuous piecewise-linear x-axis mapping. **c,** Time-series line plot of size fraction percentages at the above time points. (i) Percentage curves over the full-time range of days 0–1230; (ii) expanded view of days 0–67. Fluctuations in size-fraction proportions were pronounced: particles above 590 nm decrease markedly, whereas particles below 590 nm increase markedly; after the CCD time point, the fractions smaller than 590 nm came to occupy a slight majority, with the largest fluctuations occurring in the 80–300 nm and 300–590 nm fractions. While total particle counts fluctuated only slightly (b), panel (c) shows that the proportion of large particles ($\varphi > 590$ nm) declined sharply—from 93.86% down to 56.98% at the CCD point—while the percentages of the two smaller size fractions rose gradually amid minor fluctuations, with maximum increases in the 80 nm $< \varphi \leq 300$ nm and 300 nm $< \varphi \leq 590$ nm ranges from 2.41% to 50.23% and from 3.74% to 72.58%, respectively.
